# Acute antidepressant effect of ayahuasca in juvenile non-human primate model of depression

**DOI:** 10.1101/254268

**Authors:** Flávia Santos da Silva, Erick Allan dos Santos Silva, Geovan Menezes de Sousa Junior, Joao Paulo Maia-de-Oliveira, Vanessa de Paula Soares Rachetti, Draulio Barros de Araujo, Maria Bernardete Cordeiro de Sousa, Bruno Lobão Soares, Nicole Leite Galvão-Coelho

**Affiliations:** Postgraduate Program in Psychobiology, Federal University of Rio Grande do Norte, Natal, RN, Brazil; Laboratory of Hormone Measurement, Department of Physiology, Federal University of Rio Grande do Norte, Natal, RN, Brazil; Department of Clinical Medicine, Federal University of Rio Grande do Norte, Natal, RN, Brazil; National Institute of Science and Technology in Translational Medicine Natal, RN, Brazil; Department of Biophysics and Pharmacology, Federal University of Rio Grande do Norte, Natal, RN, Brazil; Brain Institute, Federal University of Rio Grande do Norte, Natal, RN, Brazil; Department of Physiology, Federal University of Rio Grande do Norte, Natal, RN, Brazil.

**Keywords:** translational animal model, non-human primate, common marmoset, behaviors, cortisol, early-age depression, psychedelic drugs

## Abstract

The incidence of major depression in adolescents, aged between 15 to 18 years, reaches approximately 14%. Usually, this disorder presents a recurrent way, without remission of symptoms even after several pharmacological treatments, persisting through adult life. Due to the relatively low efficacy of commercially available antidepressant, new pharmacological therapies are under continuous exploration. Recent evidence suggests that classic psychedelics, such as ayahuasca, produce rapid and robust antidepressant effects in treatment-resistant depression patients. In this study, we evaluated the potential of antidepressant effects of ayahuasca in a juvenile model of depression in a non-human primate, common marmoset (*Callithrix jacchus)*. The model induces depressive-like symptoms by chronic social isolation (60 days) and antidepressant effects monitoring included fecal cortisol, body weight, and behavioral parameters. The animals presented hypocortisolemia and the recovery of cortisol to baseline levels started already at 24h after the ingestion of ayahuasca, but not the vehicle. Moreover, in males, ayahuasca, and not the vehicle, reduced the scratching, a stereotypic behavior, and increased the feeding. Ayahuasca also improving body weight to baseline levels in male and female common marmosets. The behavioral response induced by ayahuasca shows long effect, lasting 14 days. Therefore, for this translational animal model of juvenile depression, it could be proposed that ayahuasca treatment presented more notable antidepressant effects than tricyclic antidepressant nortriptyline, investigated by our group, using this same protocol in an anterior study. Ayahuasca produced faster and more durable effect on reversion of physiological changes and depressive-like symptoms. Therefore, the results found for ayahuasca treatment corroborates in the validation of this substance as an effective antidepressant drug and encourages the return of studies with psychedelic drugs in the treatment of humor disorders, including adolescents with early-age depression.

## INTRODUCTION

Major depressive disorder (MDD) is characterized by depressed mood, anhedonia, weight alterations (loss or weight gain), sleep disorders (insomnia or hypersomnia) and psychomotor alterations (motor retardation or agitation) [Diagnostic and Statistical Manual of Mental Disorders 5, 1, 2]. Depression is currently ranked by World Health Organization (WHO) as the major contributor to global disability and suicidal deaths [3].

MDD has been associated with physiological dysregulation and a consistent finding is related to the monoaminergic unbalance, where decreased levels in dopaminergic, noradrenergic and serotoninergic neurotransmission pathway have been frequently observed [4]. However, antidepressants drugs that target these systems and increase the levels of monoamines in synaptic cleft have failed to revert depressive symptoms in totality [5] and only around 50% of patients achieve remission of symptoms after several treatments [6]. Moreover, commercially available antidepressants usually take around two weeks to achieve the desired therapeutic effects [8, 9]. Therefore, enormous efforts have been devoted to the search for alternative pharmacological treatments that improve efficacy, with a faster onset of therapeutic effects [10].

Ayahuasca is a decoction made from a combination of two plants from Amazon rainforest: *Psychotria viridis* and *Banisteriopsis caapi* [11]. Recent studies suggest that ayahuasca does not exhibit tolerance and is not addictive [12, 13, 14, 15]. Different ayahuasca compounds act on biological systems involved in the etiology of depression. For instance, N,N-dimethyltryptamine (N,N-DMT) besides acting as a serotonergic (5-HT2A) agonist [16, 17, 18] also modulates sigma-1 receptors (σ1R) that, in turn, has recently been implicated in depression [19, 20, 21]. When consumed orally DMT is inactivated by monoamine oxidase (MAO) present in the intestine and liver [22]. Ayahuasca also contains β-carbolines (Tetrahydroharmine, Harmine and Harmaline) that act as reversible monoamine oxidase inhibitors (MAOi), allowing monoamines to remain more time within the synaptic cleft space, and protecting DMT against degradation when ingested [23].

Besides MAOi properties, THH is also a serotonin reuptake inhibitor [24] and recent studies suggest significant antidepressant effects of Harmine, in a rodent model of depression [25]. Furthermore, it has been observed that ayahuasca modulates plasma cortisol, which is also involved in etiology of depression disorders, and patients with major depression show consistent altered levels in plasma and saliva cortisol [26, 27, 28] whereas healthy volunteers show increased cortisol levels 2 hours after ayahuasca intake [29, 23].

In fact, positive health benefits have been found in regular ayahuasca users in religious contexts [30, 15, 31]. Furthermore, the antidepressant effects of ayahuasca have been explored and a recent randomized controlled trial suggested a fast onset of antidepressant effect in patients with treatment-resistant depression [32, 9, 33].

Adolescents, aged between 15 to 18 years, have shown incidence rates of depression that reach to 14%, with approximately 40% of recurrence in the next 3 to 5 years following the first episode [34]. The influence of sexual steroids at this age opens an important biological window of plasticity in the nervous system, which turns the brain largely susceptible to environmental influences. If the stimulus induces maladaptive changes in brain morphology and functions, these can lead to permanent impairment of cognitive, behavioral and physiological mechanisms, increasing the probability of the emergence of mood disorders that can persist into adulthood [35, 36, 37]. Therefore, the rise of sexual hormones at puberty turns adolescents into a higher risk group of presenting depressive episodes. According to recent findings, a fast antidepressant action of ayahuasca in adult patients with treatment-resistant depression has been reported [32, 9, 33]. Thus, a question that arises from such findings is whether ayahuasca might be effective to other age ranges also susceptible to depression such as adolescents.

Therefore, this study evaluated the acute antidepressant effects of ayahuasca on physiological (cortisol fecal), body weight and behavioral parameters in a juvenile model of depression, common marmoset (*Callithrix jacchus*), after induction of a depressive-like state by chronic social isolation.

## METHOD

### Animal maintenance

All animals were housed according to the guidelines of the Brazilian Institute of Environment and Renewable Natural Resources (IBAMA) (Normative Instruction no. 169 of February 20, 2008), and care standards for animals established by the National Council for Animal Experimentation Control (CONCEA), Law No. 11.794, (October 8, 2008). In addition, the laboratory complies with international standards for *ex situ* maintenance of animals as defined by the Animal Behavior Society and the International Primatological Society. The study and experimental procedures were approved by the Animal Research Ethics Committee, (UFRN protocol No. 034/2014).

To allow genetic variability, all juvenile animals (7 to 9 months) used in this study (n = 15; 8 males and 7 females) were randomly selected from 10 different families from approximately 150 marmosets living in captive conditions in the Laboratory of Advanced Studies in Primates, at the Federal University of Rio Grande do Norte (UFRN), Natal, Brazil. The marmoset colony is formed by animals that were born in captivity as well as those captured from nature and introduced in the colony.

At baseline conditions, animals were living with their families in masonry cages (2.0 × 2.0 × 1.0 m^3^), located outdoors, under natural conditions of lighting, humidity, and temperature. The cage has on the front a glass wall with a unidirectional visor and on the back wall, a wire mesh door where water bottle and food plate are available. Inside the cage there is a nest box for resting, planks of wood, and branches of plants for environmental enrichment, allowing the animal´s displacement within the cage.

The model of depression was induced by social isolation [38]. During this procedure, animals were moved from their family groups and placed alone in a new smaller masonry outdoor cage (1.0 × 2.0 × 1.0 m^3^), but without space restrictions. During this condition, animals did not have any visual contact with related conspecifics but had auditory and olfactory contact with other conspecifics living at the colony.

None of the animals had been used in previous scientific study neither separated from their respective family groups for prolonged periods. All animals were habituated to the presence of the researchers prior to the study. Veterinary care was provided throughout the experiment. Water was available without restriction during the entire study and all animals were fed the same diet, twice per day, which included seasonal fruits such as banana, papaya, melon, and mango, as well as potato and a protein potage containing milk, oats, egg, and bread. Twice a week, a multivitamin supplement (Glicocan) was diluted in the food. The animals were weighed every 15 days, in order to monitor animal health.

### Study design

A previous study has validated the induction of depression-like state for juvenile common marmoset [38], and the same design was used herein.

During *baseline* phase (BL), juvenile marmosets (8 males and 7 females) were observed for 4 weeks while living within their families, which were followed by an eight weeks period of *social isolated context* (IC). Subsequently, 5 males and 4 females were randomly selected to be treated, first with one administration of vehicle containing a pure saline, *vehicle treatment* phase (VE), which lasts one week and comprised behavioral and physiological sampling. In the following phase, *pharmacological treatments* (PH), that also lasts one week, animals received one dose of the ayahuasca and were sampled daily. PH was followed by one more week of sampling that corresponded to *tardive-pharmacological effects* (tPE) phase. Figure 1 shows the experimental design. Besides behaviors and fecal cortisol were monitoring, body weight was also recorded during all five phases of the study. For more detailed description of the protocols, see Galvão-Coelho *et al*. [38].

Differently of traditional antidepressants that are administered usually daily, ayahuasca treatment consisted in a single dose, considering the observation that in previous studies ayahuasca provoked acute increases in cortisol levels in plasma [32], we decided to investigate behavioral and physiological parameters 24 and 48 hours after administration, day 1 (D1) and day 2 (D2) respectively, for both treatments, vehicle (VE) and ayahuasca (PH).

**Figure 1.**
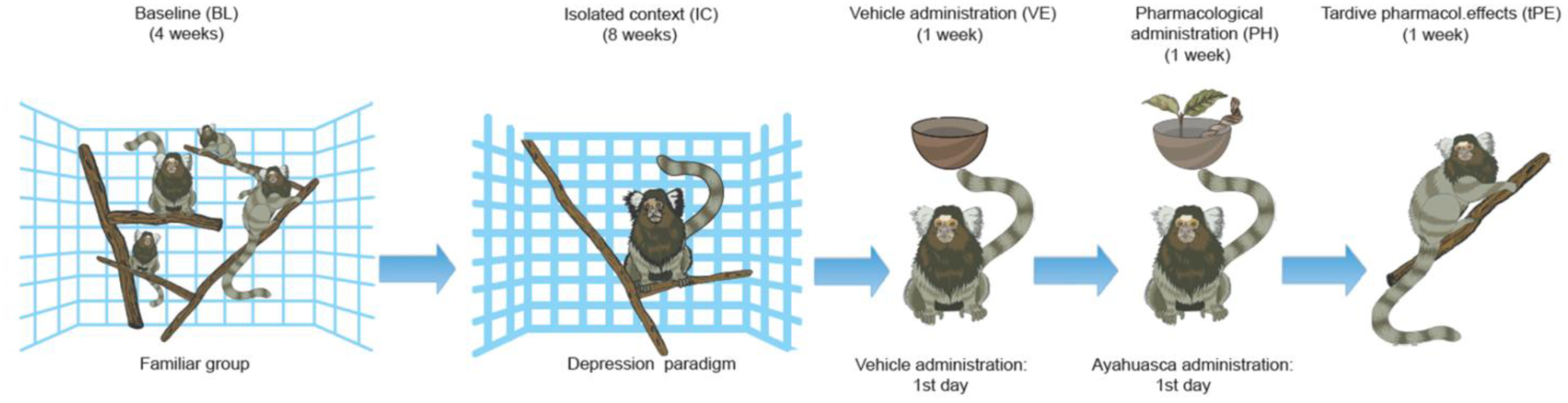
Experimental design, comprising baseline (BL, 4 weeks), social isolated context (IC, 8 weeks), vehicle treatment (VE, 1 week), pharmacological treatments (PH, 1 week) and tardive-pharmacological effects (tPE, 1 week) phases, where marmosets were sampled for behaviors, fecal cortisol and body weight.

### Treatments

All animals were treated with one administration of the vehicle, and seven days after, the same animals received a single dose of ayahuasca. In both cases, we used a dose of 1.67 ml/ 300g of animal weight, via gavage. The dose of ayahuasca established for marmosets was transposed from the dose used in humans, using a recommended and effective allometry procedure [39].

The ayahuasca used was produced and provided free of charge by the religious organization called “Barquinha” (Brazil). A unique ayahuasca batch was used throughout the whole study, for all animals. The preparation of the tea followed a traditional recipe, by infusing 50% of leaves of *Psicotria viridis* with 50% of the stem of *Bansteriopsis caapi*. The batch was stored in bottles in a refrigerator. The quantification of alkaloids was determined by mass spectroscopy by the Laboratory of Toxicological Analysis at the University of São Paulo. Results showed 0.36 ± 0.01mg of DMT/mL, 1.86 ± 0.11mg of Harmine/mL, 0.24 ± 0.03mg of Harmaline/ml of and 0.20 ± 0.05mg of tetrahydroharmine (THH)/mL [40].

### Behavioral recordings

The recorded behaviors were the same validated by Galvão-Coelho *et al*. [38] for juvenile marmosets, and included: species-specific behaviors, as scent marking (frequency), individual piloerection (frequency), scratching (duration), autogrooming (duration), and behaviors associated with depressive-like state that can be compared across species such as locomotion (frequency), somnolence (duration), feeding (duration) and anhedonia. Anhedonia was measured by frequency and duration of ingestion of an aqueous solution of sucrose (4.16%). For more detailed description of behavioral data and its implications in stress context, see Galvão-Coelho *et al*. [38].

The selected behaviors were recorded between 6:30 and 7:30 a.m. to avoid the influence of circadian variation, over a 30-min period by the focal continuous method [41].

### Fecal collection and cortisol assay

Fecal samples were collected between 6:30 and 8:30 a.m. to avoid circadian variation in cortisol profile in feces [42]. Cages were cleaned prior to fecal collection, to avoid collecting samples expelled prior to 6:30 a.m. Samples were stored at -4 °C until cortisol extraction and quantification, which was measured at the Hormonal Measurements Laboratory of the Department of Physiology (UFRN), according to the protocol of Sousa and Ziegler [43]. Fecal cortisol reflects plasma cortisol with a delay of approximately 8–10 hours. Intra- and inter-assay coefficients of variation were 2.74% and 16.61%, respectively.

### Statistical analysis

Hormonal data were normalized by logarithmic transformation and for both hormonal and behavioral data, the statistical technique of bootstrap resampling was applied to the multivariable analysis. General linear model (GLM) and Fisher’s *post hoc* tests were used to investigate the variations of behaviors, cortisol, and body weight, between sex, throughout the study phases. Additionally, the parametric Student’s *t*-test was used to analyze hormonal and behavioral data between specific phases. Statistically significant results were considered for p ≤ 0.05.

## RESULTS

The analysis of studied variables showed at the end of IC, a significant decrease in cortisol levels, increase in autogrooming, scratching and somnolence, as well as reductions in feeding and sucrose ingestion. All these changes were independent of sex. (Table 1).

**Table 1.** Statistical values, GLM test and LSD *post-hoc*, and direction of alterations of physiologic and behavior parameters in response to social isolated context, compared to baseline.

| VARIABLES | STATISTICAL VALUES | ALTERATION |
| --- | --- | --- |
| **Cortisol** | F= 4.38 p= 0.03 df= 1<sup>P</sup><br>F= 0.00 p=0.98 df= 1<sup>P/S</sup> | ↓<br>--- |
| **Autogrooming** | F= 15.31 p= 0.00 df= 1<sup>P</sup><br>F= 0.42 p= 0.73 df= 1<sup>P/S</sup> | ↑<br>--- |
| **Somnolence** | F= 3.73 p= 0.05 df= 1<sup>P</sup><br>F= 1.47 p= 0.22 df= 1<sup>P/S</sup> | ↑<br>--- |
| **Feeding** | F= 18.10 p= 0.00 df= 1<sup>P</sup><br>F= 0.90 p= 0.34 df= 1<sup>P/S</sup> | ↓<br>--- |
| **Ingestion of sucrose**<br>(frequency) | F= 22.61 p= 0.00 df= 1<sup>P</sup><br>F= 5.69 p= 1.16 df= 1<sup>P/S</sup> | ↓<br>--- |
| (duration) | F= 5.55 p= 0.02 df= 1<sup>P</sup><br>F= 0.12 p= 0.72 df= 1<sup>P/S</sup> | ↓<br>--- |
| **Scratching** | F=4.72 p= 0.03 df= 1<sup>P</sup><br>F=30.14 p= 0.00 df= 1<sup>P/S</sup> |  |
| Male | p= 0.00 | ↑ |
| Female | p= 0.02 | ↑ |
| **Scent-marking** | F= 5.49 p= 0.02 df= 1<sup>P</sup><br>F= 9.68 p= 0.00 df= 1<sup>P/S</sup> |  |
| Male | p= 0.00 | ↑ |
| Female | p= 0.15 | --- |
| **Body weight** | F= 9.12 p= 0.00 df= 1<sup>P</sup><br>F= 11.27 p= 0.00 df= 1<sup>P/S</sup> |  |
| Male | p= 0.00 | ↓ |
| Female | p= 0.81 | --- |
| **Locomotions** | F= 0.03 p= 0.85 df= 1<sup>P</sup><br>F= 0.32 p= 0.57 df= 1<sup>P/S</sup> | ---<br>--- |
| **Individual piloerection** | F= 0.03 p= 0.08 df= 1<sup>P</sup><br>F= 0.07 p= 0.78 df= 1<sup>P/S</sup> | ---<br>--- |

Acronyms: <sup>P</sup> = statistical analyze for phase, <sup>P/S</sup> = statistical analyze for interaction between phase and sex.

For scent marking and body weight, sexual dimorphic profiles of variation were observed, where only males showed increased scent marking and reduced body weight (Table 1). No significant statistical variations were observed in locomotion and individual piloerection with social isolation (Table 1).

After ayahuasca treatment (PH), but not after vehicle (VE), males decreased scratching with respect to the IC. Such reduction in the PH lasted more 7 days, being also observed in the tPE (Figure 2A and Table 2). Again, only males showed an increase in feeding after treatment with ayahuasca, which was sustained until tPE, and did not vary after treatment with vehicle (Figure 2B and Table 2). Both sexes increased body weight after ayahuasca, but not after vehicle, and was sustained throughout tPE (Figure 2C and Table 2). Body weight gain, induced by ayahuasca, allowed the recovery to baseline weight levels (PT: t= -1.68, p=0.09 / tPE: t = -1.33, p = 0.18).

**Figure 2.**
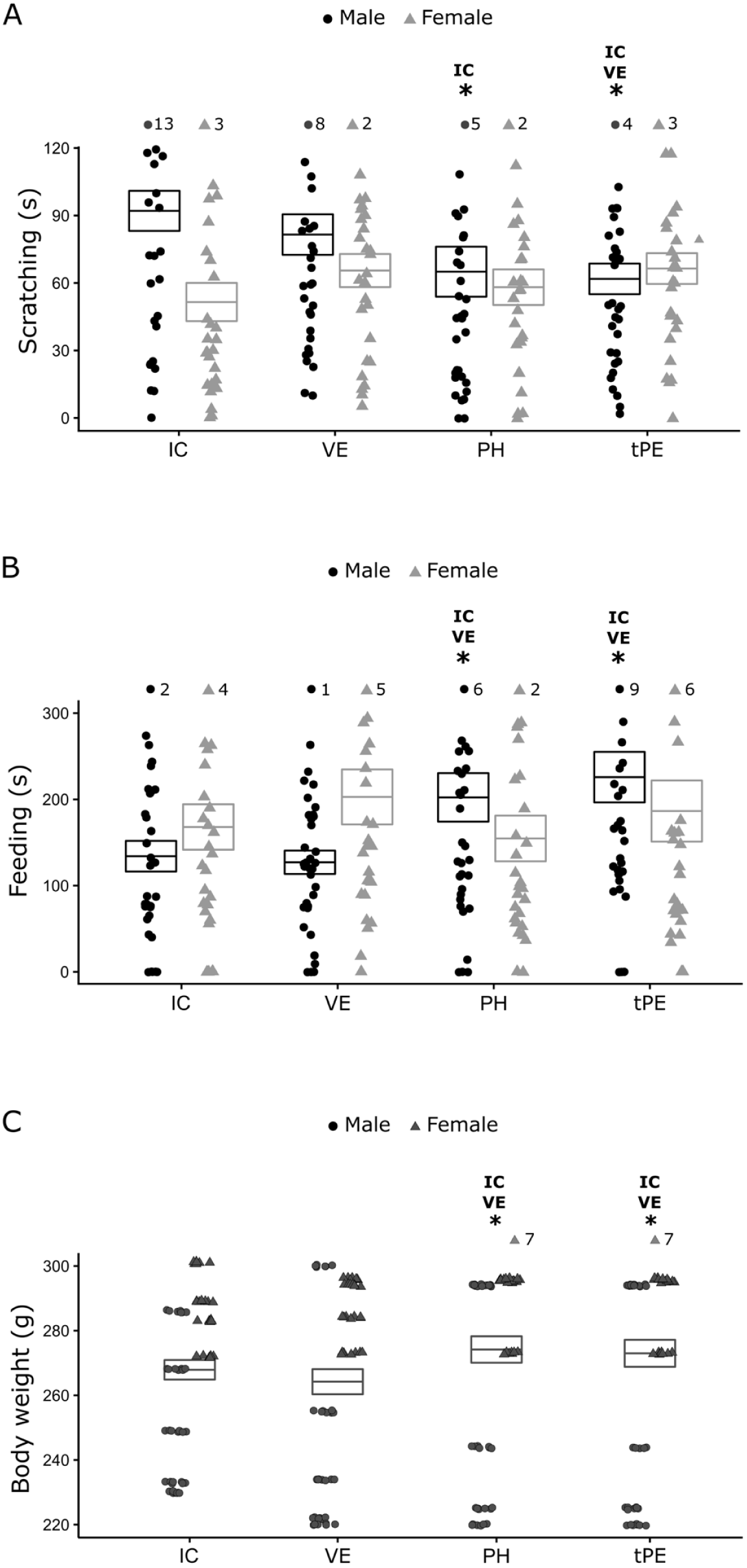
Means ± SEM of behaviors: (A) scratching, (B) feeding, (C) body weight in male and female juveniles *C. jacchus* along IC= isolated context, VE = Vehicle treatment, PH = Pharmacological treatment and tPE = tardive-Pharmacological Effects. * = statistically significant difference between respective phase and phase indicated (s) next to the symbol. GLM test and *post hoc* Fisher, p ≤ 0.05.

**Table 2.** Statistical values, GLM test and LSD *post-hoc,* and direction of alterations of physiologic and behavior parameters in response to pharmacological treatments, compared with isolated context.

| VARIABLES | STATISTICAL VALUES | VE | PT | tPE |
| --- | --- | --- | --- | --- |
| **Cortisol** | F= 1.18 p=0.31 df = 3<sup>P</sup><br>F= 0.42 p=0.37 df = 3<sup>P/S</sup> | --- | --- | --- |
| **Autogrooming** | F= 0.93 p=0.42 df = 3<sup>P</sup><br>F= 1.45 p=0.22 df = 3<sup>P/S</sup> | --- | --- | --- |
| **Scratching** | F= 1.31 p=0.27 df = 3<sup>P</sup><br>F= 3.59 p=0.01 df = 3<sup>P/S</sup> |  |  |  |
| *Male* |  | p= 0.27 | p= 0.00 ↓ | p= 0.00 ↓ |
| *Female* |  | p= 0.19 | p= 0.53 | p= 0.16 |
| **Somnolence** | F= 0.57 p=0.63 df = 3<sup>P</sup><br>F= 0.77 p=0.51 df = 3<sup>P/S</sup> | --- | --- | --- |
| **Feeding** | F=2.23 p=0.08 df = 3<sup>P</sup><br>F = 3.55, p = 0.01, df = 3<sup>P/S</sup> |  |  |  |
| *Male* |  | p=0.81 | p=0.02 ↑ | p=0.00 ↑ |
| *Female* |  | p=0.29 | p=0.69 | p=0.57 |
| **Ingestion of sucrose (frequency)** | F= 13.28 p=0.80 df = 3<sup>P</sup><br>F= 0.78 p=0.50 df = 3<sup>P/S</sup> | --- | --- | --- |
| **Scent-marking** | F= 3.14 p=0.06 df = 3<sup>P</sup><br>F= 5.30 p=0.18 df = 3<sup>P/S</sup> | --- | --- | --- |
| **Body weight** | F = 15.35, p= 0.00, df = 3<sup>P</sup><br>F= 2.00 p=0.11 df = 3<sup>P/S</sup> | p=0.02 ↓ | p=0.00↑ | p=0.00↑ |
| **Locomotion** | F= 1.01 p=0.38 df = 3<sup>P</sup><br>F= 1.80 p=0.14 df = 3<sup>P/S</sup> | --- | --- | --- |

Acronyms: <sup>P</sup> = statistical analyze for phase, <sup>P/S</sup> = statistical analyze for interaction between phase and sex.

No significant alterations in response to both treatments, vehicle and ayahuasca, were observed in fecal cortisol (Figure 3A and Table 2), as well as in autogrooming, scent marking, locomotion, ingestion of sucrose and somnolence (Table 2). With respect to individual piloerection, it was not possible to perform GLM analyzes due to the large number of zero frequencies and low variation of data.

**Figure 3.**
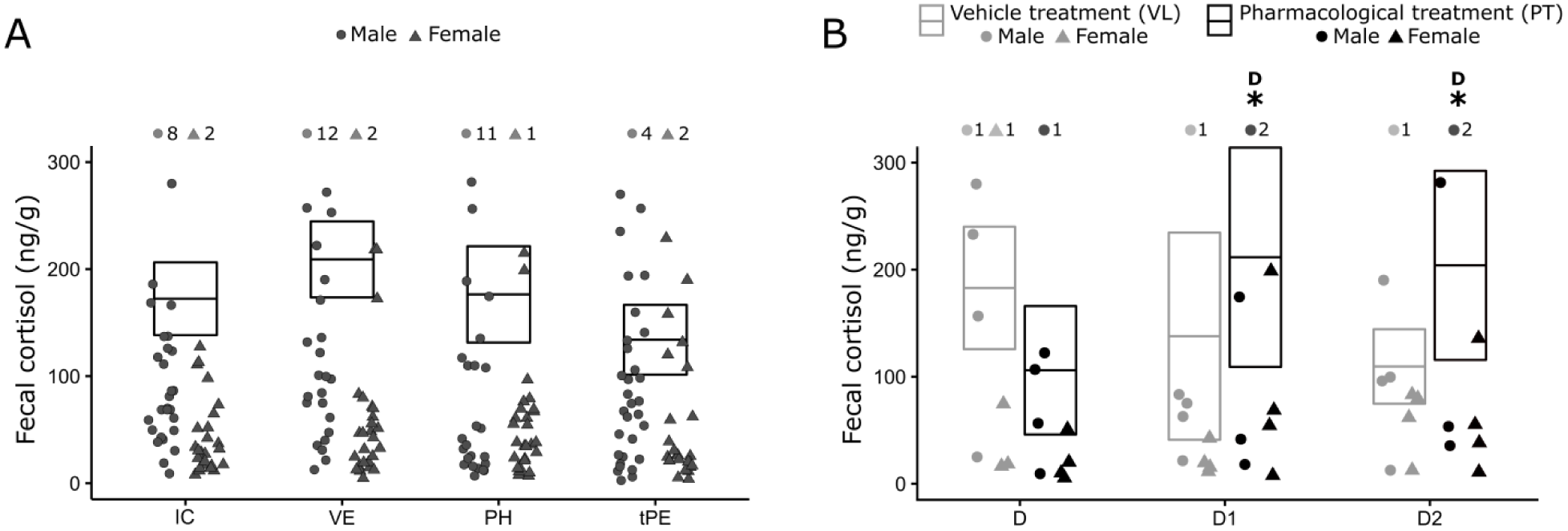
Means ± SEM of fecal cortisol: A) along IC= isolated context, VE = Vehicle treatment, PH = Pharmacological treatment and tPE = tardive-Pharmacological Effects, and B) At D1 (24h) and D2 (48h) after treatment with Vehicle (VE) and ayahuasca = PH. * = statistically significant difference between respective phase and phase (s) indicated (s) next to the symbol. GLM test and *post hoc* Fisher, p< 0.05.

Cortisol levels increased 24 hours (D1) and 48 hours (D2) after ayahuasca ingestion, but not after vehicle, (Figure 3B and Table 3) Moreover, cortisol levels observed at D1 and D2 returned to similar values found in BL (D1: t = -0.55, p=0.58 / D2: t= -1.30, p= 0.21). No significant alterations were found in all behaviors at D1 and D2 in response to vehicle or ayahuasca (Table3). The GLM analysis of individual piloerection and sucrose ingestion carried out due a large number zero’s and the low variability of the data.

**Table 3.**
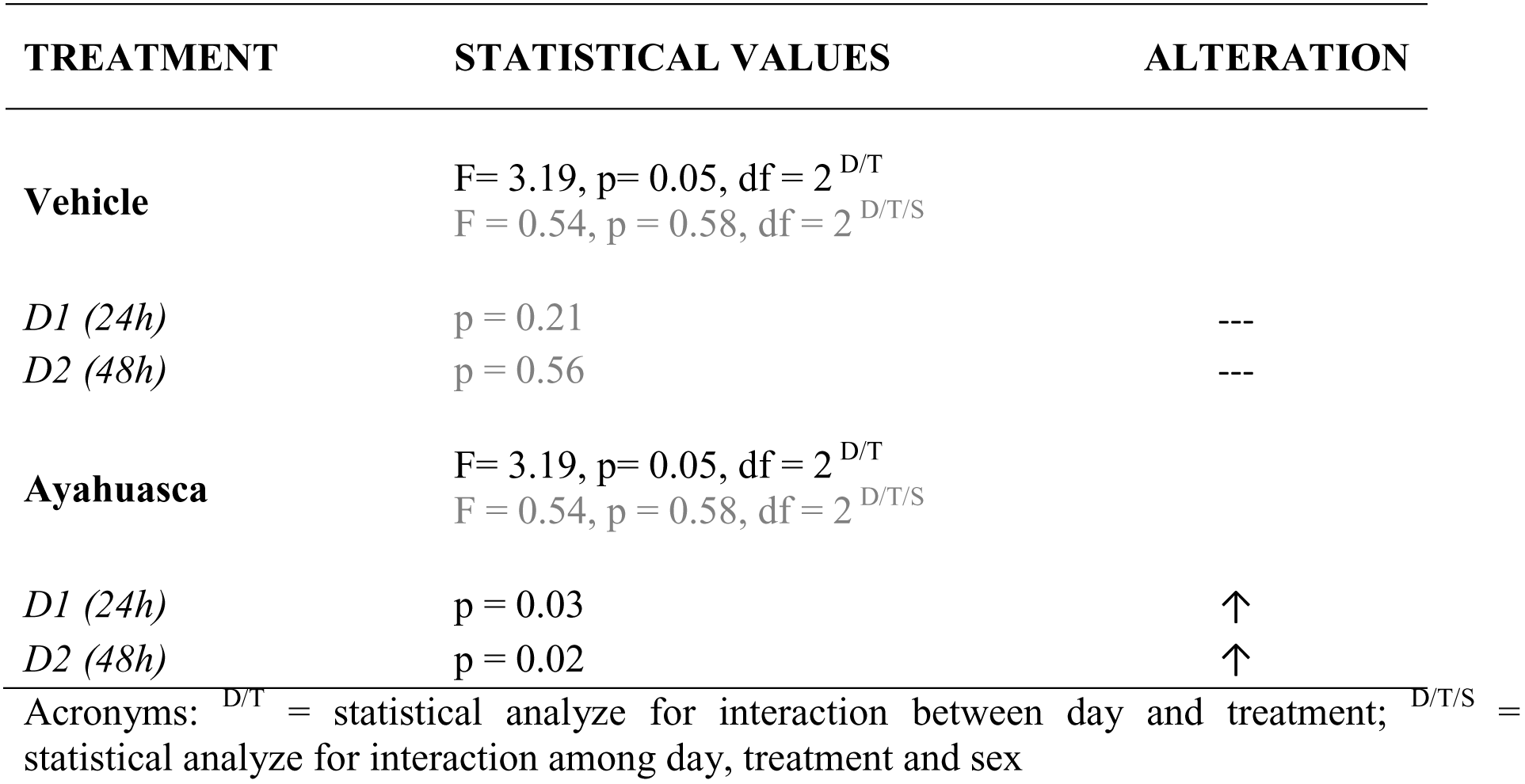
Statistical values, GLM test and LSD *post-hoc,* and direction of alterations of acute cortisol in response to pharmacological treatments, in D1 (24h) and D2 (48h) after treatment with Vehicle (VE) and ayahuasca (PH).

| TREATMENT | STATISTICAL VALUES | ALTERATION |
| --- | --- | --- |
| **Vehicle** | $F = 3.19, p = 0.05, df = 2^{D/T}$<br>$F = 0.54, p = 0.58, df = 2^{D/T/S}$ | |
| D1 (24h) | $p = 0.21$ | --- |
| D2 (48h) | $p = 0.56$ | --- |
| **Ayahuasca** | $F = 3.19, p = 0.05, df = 2^{D/T}$<br>$F = 0.54, p = 0.58, df = 2^{D/T/S}$ | |
| D1 (24h) | $p = 0.03$ | ↑ |
| D2 (48h) | $p = 0.02$ | ↑ |

Acronyms: $D/T$ = statistical analyze for interaction between day and treatment; $D/T/S$ = statistical analyze for interaction among day, treatment and sex

## DISCUSSION

This study found a rapid antidepressant effect of ayahuasca on behavioral expression in males and females juvenile common marmosets presenting depressive-like state induced by chronic social isolation. In addition, both fecal cortisol and body weight returned to baseline levels, when the animals were living in their family groups. Moreover, some behavioral alterations indicative of depression-like state was reduced mainly in males.

After being moved from the family group and remain socially isolated during eight weeks (IC), marmosets increase auto-direct stereotypic behaviors as scratching and autogrooming. In non-human primates such behaviors have been expressed during psychosocial stress [44, 45]. Feeding reduction, increases in somnolence and anhedonia, inferred here by reduction sucrose ingestion, for both sexes were also observed after social isolation. Males also showed body weight loss and increase in scent marking, which in context of stress is considered as an anxious behavior, because it occurs without a defined interest, such as those involved in territorial defense and reproductive signalization [46]. The presence of anhedonia, somnolence, reduction in feeding and body weight changes are consistent with symptoms observed in patients with depression and are used as guidelines to perform the diagnostic of this pathology as described in diagnostic and Statistical Manual of Mental Disorders (DSM-5).

After treatment with the vehicle, no changes were recorded for any behaviors or body weight. However, a single dose of ayahuasca improved some of these symptoms of the depressive-like state, mainly in males. In this case, a significant reduction of scratching and increased feeding indicates a positive effect on the recovery of such as functions. Although in females no changes in feeding have been observed, body weight regulation to baseline levels occurs for both sexes. In a previous study where the antidepressant nortriptyline was used, in similar protocol and animal model, only females showed reductions in scratching after treatment [34]. The different action of ayahuasca and nortriptyline to improve depressive symptoms in male and female common marmosets points out to the importance of studying both sexes in translational studies of depression. Furthermore, despite in humans sex difference in response to antidepressants has been recorded, males of animal models continue to be more frequently used in experimental protocols for depression studies [47, 48].

The positive ayahuasca modulation observed in feeding behavior is potentially important for patients with depression with loss of appetite and body weight. The anorectic effect of tricyclic antidepressants observed by Galvão-Coelho et al. [34] after treatment with nortriptyline in common marmosets is also perceived in patients with depression and it is considered a side-effect that induces a minor tolerance.

Available studies with ayahuasca and animal models of depression until the present moment did not use non-human primates as animal models and also did not use juvenile animals, this is the first one. Normally, the studies use adult rodents as animal models of depression [25, 49], species phylogenetically more distant of humans than common marmosets. Despite these studies also observed positive antidepressant effects with the use of ayahuasca, or it´s specific components, differently of the present study, they used 14 days of treatment and did not observe the continued response after treatment stopped. For instance, rats treated with Harmine (5, 10 and 15 mg/kg) for 14 days showed improvements in forced swimming and open-field tests [25]. Wistar female rats treated with ayahuasca for 14 days also presented better performance in forced swimming test, when compared to a group treated with fluoxetine [49]. On the other side, this study showing improvement in body weight and depressed-like behaviors that remained until 14 days after one single-dose ayahuasca treatment.

Besides of the well-known pharmacological action of ayahuasca, such as antagonist of MAO and the transporter of serotonin and agonist of the 5-HT-2 receptor, others pharmacological targets can be involved in this rapid antidepressant effect of ayahuasca observed in juvenile’s marmosets. Recently, some antidepressant effects in rodents, as a reduction of anhedonia, were associated with Sigma-1 receptors activation [20, 21]. Moreover, some indirect pathways modulate by DMT agonist action on 5-HT2a and sigma-1 receptors can stimulate molecular and cellular events involved in neural and synaptic plasticity, such as encoding of transcription factors (c-fos, egr1, egr-2), synthesis of brain-derived neurotrophic factor (BDNF),enhances of CaMKII /CaMKIV and protein kinase B (Akt) activities in hippocampus, all compatible with antidepressant action [50, 51, 20].

Ayahuasca treatment did not induced alterations in autogrooming, scent marking, somnolence and ingestion of sucrose solution similar to that results observed after treatment with nortriptyline [34]. The absence of drug modulation in these behaviors might be related with the dose and duration of ayahuasca treatment, which might be not enough to promote a stronger antidepressant effect, suggesting that alternatives protocols should be tested in order to verify a more robust behavioral improvement in juveniles’ marmosets.

Regarding cortisol variation across the study’ phases, after chronic social isolation of eight weeks, common marmosets showed significantly low levels of fecal cortisol. Low levels of cortisol have been reported after strong stressors both in humans and in small animals [52, 53, 54]. For instance, in juvenile common marmosets exposed to the repeated separation of their families in infancy [55] or chronic social isolation along the juvenile stage [56, 34]. A recent study with common marmosets found that 21 days of social isolation in juvenile stage is enough to reduced cortisol to levels below baseline [56]. Hypocortisolemia also was correlated to depression-like state in adult females of *Macaca fascicularis* [54]. Moreover, previous studies have reported hypocortisolemia in patients with atypical unipolar major depression and major depression with remittent conditions [57, 58, 33].

During prolonged stress response, the complex systems of the interaction of negative feedbacks of hypothalamus-pituitary-adrenal (HPA) axis could turn imbalanced and changes adrenal function, which in turns reduces cortisol synthesis [59]. A sustention of cortisol at low levels deregulates all system of adaptation since cortisol is a pleiotropic hormone that regulates hormonal, neural and immune system responses to challenging situations [60, 61]. Individuals that show a chronic decrease in cortisol levels normally present weakness, weight loss and immunological dysfunction [62].

Low levels of cortisol observed after isolation started rising already at 24 and 48 after treatment with ayahuasca, recovering cortisol to baseline levels. The observed homeostatic regulation was fast and did not extend into the later phases (PT and tPE). A previous study with the same animal model of juvenile depression, but treated with nortriptyline chloride (Pamelor^TM^), revealed that nortriptyline increases cortisol occurred after one week, and the raise overpassed baseline levels. These suggest that ayahuasca induced faster and more adjusted regulation in cortisol levels than nortriptyline. Furthermore, previous studies with ayahuasca have been suggesting antidepressant effects of ayahuasca in a treatment resistant depression already one day after a single dosing session with ayahuasca [9, 32]. It is interesting, however, to note that the rapid antidepressant effect observed in both cases of use of ayahuasca, classic antidepressants usually take at least two weeks to reach the desired therapeutic response [63, 64].

The effects of antidepressants on the HPA axis depend on the class of the drug (MAOi, tricycle, Selective Serotonin Reuptake Inhibitors, or others) [65, 66]. Serotoninergic agonists drugs, such as ayahuasca and nortriptyline, might modulate both secretion of corticotropin-releasing hormone (CRH) and/or adrenocorticotropic hormone (ACTH), at the hypothalamic and pituitary gland, respectively [67, 68]. Moreover, the duration of treatment also is an important issue in HPA axis modulation. Normally, acute treatment increases cortisol levels and long-term ones, in the opposite, induces a reduction. Probably, reduced cortisol secretion by adrenals is due to an up-regulation of glucocorticoids receptors (GR and MR) in the brain, which in turn can increase negative feedback [69, 70].

The differences in the modulation of cortisol levels by ayahuasca and nortriptyline might be in part due to its distinct chemical proprieties and duration of treatment. Ayahuasca was administrated only once to animals, whereas in our previous study nortriptyline was injected during seven consecutive days [34]. Some anterior studies have proposed that a single dose of ayahuasca in humans is enough to improve clinical symptoms for seven days [9, 32] and to regulate salivary and plasmatic cortisol to homeostatic levels [33].

In summary, behavioral symptoms, body weight and cortisol profiles of depression found in male and female juvenile common marmosets exposed to chronic social isolation (8 weeks) were large part reverted by ayahuasca treatment, more prominently in males. Nevertheless, when the effectiveness of ayahuasca was compared with nortriptyline, ayahuasca, apparently, showed more remarkable antidepressant results than nortriptyline, since the start of improvements of physiological alterations and behavioral symptoms of depression were faster, long lasting and adjusted.

Therefore, this study carries significant additional evidences that support the antidepressant action of ayahuasca using a depression animal model phylogenetically more closely of humans than rodents, a non-human primate. Moreover, for the first time, the therapeutic value of ayahuasca as an effective antidepressant drug in juvenile age was demonstrated. All anterior studies with ayahuasca were developed with adult animal models or human, this is the first study with juveniles. As puberty is an important ontogenetic period of brain plasticity, a treatment presenting fast antidepressant effects, emerges for the first time, to our knowledge, as a good alternative. Further studies may evaluate the safety and tolerability of the use of ayahuasca as an antidepressant treatment at a young age.

## Conflict of Interest Statement

This research was conducted in the absence of any commercial or financial relationships that could be construed as a potential conflict of interest.

## Author and Contributors

Designedtheexperiment:Galvão-CoelhoN.L.,Maia-de-OliveiraJ.P,DeAraujoD.B.,

Soares, B.L, Rachetti V.P.S. and Sousa M.B.C.;

Collected experimental data: Da Silva F. S.; Silva, E. A. S.; Sousa Junior, G. M.;

Carried out statistical analysis: Da Silva F. S.;

Arranged figures: Da Silva F. S., Silva E. A. S and Soares, B.L.;

Prepared manuscript: Da Silva F.S., Galvão-Coelho N.L., SousaM. B.C.,Soares B.L, De Araujo D.B., Rachetti V.P.S. and Maia-de-Oliveira J.P.

## Acknowledgments

We would like to thank Edinolia Câmera, Antonio B. da Silva, Geniberto C. dos Santos and Janaína Nitta for animal and veterinary care and Raíssa Nóbrega de Almeida for hormonal measurements.

## Abbreviations

BL: Baseline
IC: Isolated context
VE: Vehicle Treatment
PH: Pharmacological treatment
tPE: tardive-Pharmacological Effects
D1: Day 1
D2: Day 2.

